# A Differential Ion Mobility Acoustic Ejection Mass Spectrometer System for Screening Isomerization-Mediating Enzyme Drug Targets

**DOI:** 10.1101/2024.09.25.614780

**Authors:** Samad Bazargan, Patricia Dranchak, Chang Liu, James Inglese, John Janiszewski, Bradley B. Schneider, Thomas R. Covey

## Abstract

We report the first implementation of ion mobility mass spectrometry combined with an ultra-high throughput sample introduction technology for high throughput screening (HTS). The system integrates differential ion mobility (DMS) with acoustic ejection mass spectrometry (AEMS), termed DAEMS, enabling the simultaneous quantitation of structural isomers that are the substrates and products of isomerase mediated reactions in intermediary metabolism. We demonstrate this potential by comparing DAEMS to a luminescence assay for the isoform of phosphoglycerate mutase (iPGM) distinctively present in pathogens offering an opportunity as a drug target for a variety of microbial and parasite borne diseases. The metabolome consists of many structural isomers that require for separation a mobility resolving power of more than 300. Resolving powers measured in collision cross section space of 1588 and 1948 for 2- and 3-phosphoglycerate and the citrate/isocitrate isomeric pairs respectively are shown. These are the highest reported ion mobility resolving powers for molecules from the metabolome reported to date. The potential for DAEMS as a generalized screening tool is demonstrated with the separation of the substrates and products of two additional isomerases that present as potential therapeutic targets, chorismate mutase and triosephosphate isomerase. The separations are achieved at speeds compatible with the sample introduction rates of AEMS providing sufficient data points to integrate the peaks for quantitation without the use of internal standards. DMS hyphenated with acoustic sample ejection MS provides a unique solution to high throughput mass spectrometry applications where isomer and other types of separations are required.

---

Among the biochemical cycles and pathways that drive cellular metabolism some harbor enzymes that have different isoforms in pathogens versus higher forms of life offering opportunities for potential drug targets with high therapeutic specificity. Many of these metabolic enzymes are classified as isomerases, mutases, and transferases which characteristically convert substrates into products that are structural isomers. Thirty percent of the Mass Spectrometry Metabolite Library of Standards (MSMLS) is comprised of different combinations of isomeric species that cannot be distinguished by accurate mass measurement of their molecular ions alone^1^. For many, because of their structural similarities, the collision induced dissociation (CID) spectra are identical or nearly so such that the most abundant fragment ions useful for quantitation are the same.

In recent years there has been an increasing emphasis on developing high-throughput label free screening assays (HTS) using mass spectrometry (MS) as the detection method as well as for other applications where ultra-high throughput is required^2-6^. With mass spectrometry there is the potential to measure the concentrations of the substrates and products of the enzymatic reactions used in HTS directly while screening for inhibitors of these enzymes rather than use indirect indicators involving photometric outputs generated by coupled enzyme systems. Acoustic ejection mass spectrometry (AEMS), has demonstrated the ability to detect inhibitors and monitor the kinetics of several classes of enzymes including methyl transferases^7^, acyltransferases^8-10^, demethylases^10^, dehydrogenases^11^, proteases^12^, cyclic AMP/GMP synthase^13^, cytochrome P-450 oxygenase’s^10^, cell-based transporter enzymes^14-15^, and cell-based screens^16^. Sample throughputs ranging from 1-6 samples per second and 50,000-100,000 per day make possible the identification of inhibitors from libraries containing more than one million drug candidates in less than 2 weeks, challenging the throughputs of parallel optical detectors used as plate readers^17^.

Monitoring isomerase, mutase, and transferase reactions presents a special problem for mass spectrometry detection and quantification. In addition to the substrates and products being isomers they are simultaneously present in the reaction mixture at concentrations that can differ by greater than 3 orders of magnitude requiring separation prior to MS with sufficient resolution to reduce cross channel overlap to less than 0.1%. As demonstrated here, to be compatible with the speed of high throughput mass spectrometry (HTMS) and the quantitation requirements of HTS, these separations need to be carried out in a few tens of milliseconds.

Although chromatographic and electrophoretic systems^33^ can separate structural isomers, their speeds are not compatible with HTMS. Low field ion mobility, a gas phase separation technique based on molecular size as measured by collision cross sections (CCS)^18^, has been considered an option. A thorough study of the applicability of ion mobility mass spectrometry (IM-MS) for non-targeted metabolomic studies indicated that a significant portion of isomers in the metabolome have CCS differences of < 0.5% which requires a low field mobility resolving power of 300 or greater as measured in collision cross section space^1, 19^. Using a high resolution low field mobility system with a traveling wave enabled serpentine flight path of 13 meters, resolving powers between 200-300 were achieved using flight times up to 700 msec with transmission reduction of low mass ions (< 350 *m/z*) being encountered^20^.

Achieving resolving powers in the 200-300 range is challenging for low field ion mobility systems comprised of drift tubes of various lengths and geometries, cyclic multi-pass flight paths, and those utilizing traveling waves to surf the ions through the cells. To achieve resolutions in this range requires longer flight times with a concomitant reduction in ion transmission. Resolving powers have been reported as high as 750 using a flight time of ∼1.5 seconds with a cyclic traveling wave ion mobility analyzer^21^ and 1860 using a flight time of ∼10 seconds for m/z > 600 using a SLIM devise^22^. However, the relatively long flight times put limits on its utility where ultra-high throughput is required. With HTMS, defined as 1 sample per second or greater, these separation times do not match the duty cycle of the sample introduction system. With low field mobility and a TOF MS detector ions of all m/z can be collected in a single spectrum providing a multiplexing advantage but the speed per spectrum remains too low for this application.

The separation principles of differential ion mobility (DMS) are different from low field mobility and can provide high resolution separations with flight times less than 5 msec^23-24^. Selectivity is largely derived from the gas phase chemical clustering properties of molecular ions instead of their molecular size^25^. This approach has demonstrated the ability to separate positional isomers and diastereomers^23-24^. This capability has stimulated the use of DMS as an important tool to study the flux of metabolites through metabolic cycles under different experimental conditions resulting in valuable mechanistic insights in the field of fluxomics^26-30^.

This is the first report of combining high-throughput sample introduction with IM-MS for HTS purposes. An earlier report alluded to the potential of including DMS with AEMS for drugs of abuse testing^31^. In this present report DMS was integrated with AEMS for high-speed sample introduction, euphemistically termed DAEMS, to solve the isomer separation problem in drug discovery. Separations are demonstrated for the substrates and products of phosphoglycerate mutase, chorismate mutase, triosephosphate isomerase, and aconitase isomerase with sufficient resolution to enable quantitation over a broad dynamic range. Resolving powers of 1588 for 2 & 3 phosphoglycerate and 1948 for the citrate/ isocitrate pair are shown using literature reported CCS values^1, 32^. These separations are achieved using measurement times that are compatible with ultra-high throughput AEMS.

The enzyme used to test the DAEMS concept for HTS is cofactor-independent phosphoglycerate mutase (iPGM), the isoform in pathogens of phosphoglycerate mutase which catalyzes the interconversion of 2 & 3-phosphoglycerate (2-PG; 3-PG) in the glycolytic and gluconeogenic pathways^33-35^. Because this isoform is different in various infectious organisms than mammals it presents as a potential drug target. A coupled enzyme ATP-based luminescence assay was previously developed for endpoint measurements and an NADH absorbance assay for kinetic determinations^33^. We compare the assay metrics from DAEMS to these well-established photometric based systems as a first step toward validation. The ability of DMS to resolve isomers in the metabolome at high-speed indicates the potential of DAEMS to address a broad range of HTS targets as well as other applications of HTMS requiring the separation of isomers at speeds where HPLC is not an option.

## EXPERIMENTAL SECTION

### Acoustic sample ejection module

The prototype AEMS system comprised a SCIEX 6500+ QTRAP (SCIEX, Concord, ON, Canada) equipped with an OptiFlow ESI source and a prototype acoustic sample ejection module. Details of the design and operation of the acoustic ejection sample introduction has been previously reported^36-37^. The carrier solvent pump was controlled via a LabView-based application and the acoustic ejection process was controlled by using the NECO Diagnostics software. MS data was acquired with Analyst 1.7 (SCIEX, Concord, ON, Canada) in negative ion mode. Data analysis was conducted using a SCIEX research version of the MultiQuant software package.

### Open Port Interface (OPI) module

The OPI dimensions, materials of construction, and method of tuning the transfer liquid flow rates for optimal nL droplet capture are used as detailed in previous reports^37^. To maintain solubility of 2-and 3-PG a high aqueous transport fluid of 89:10:1 H_2_O:CH_3_OH:NH_4_OH [volumetric percentage] was critical (Supplementary Info 1). An ESI emitter/transport tube having low flow resistance was used to maintain high fluid flow rates and speed as described in Supplemental Info 1. Transport fluid flow is aspirated by the constant pressure drop created by the ESI gas nebulizer so resistance to flow, and achievable flow rate, is determined by solvent viscosity and tubing dimensions.

### Differential Ion Mobility module

A breadboard Sciex Selex-ION+ DMS was utilized. Details of the design and operation of the DMS component have been previously reported^23-24^. An LC pump introduced the modifier into the curtain gas stream using a Tee configuration and via a 10cm long PEEKsil tubing with an internal diameter of 50um.

Resolving power was calculated using the collision-cross section (CCS) based relationship of McLean^20^ adapted to DMS variables. The low field mobility time-based separation variables^9^ are replaced with the compensation voltage (CoV) based separation variables in DMS. 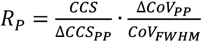 where the average CCS and CoV_FWHM_ values are used, and ΔCCS_PP_ and ΔCoV_PP_ are the differences in the collision cross section and the observed CoV peak positions, respectively. The CCS values used are from literature reported drift tube measurements^1^.

### iPGM inhibition assay

The *C. elegans* iPGM coupled-enzyme functional assay was adapted from that previously described^33^. For further details and the protocol used see Supplementary Info 8-9.

Data from both assays were normalized to DMSO neutral control and no enzyme buffer as maximum inhibition (−100% activity). Concentration response curves (CRCs) were fit in GraphPad Prism with the log(inhibitor) vs. response – variable slope (four parameters) algorithm and a constraint that absolute value of hillslope ≤3.

### Reagents and materials

See supplementary info 10 for details.

## RESULTS AND DISCUSSION

### DAEMS System Overview

*Matching Sample Introduction and Isomer Separation Speeds*. The DAEMS system is depicted in Figure 1 with a description of the relationship of each DAEMS sub-component and their basic operation provided in the figure legend. The purpose is to provide a means to execute label free mass spectrometry based HTS assays for inhibitors of metabolic targets whose substrates and products are structural isomers. An assay for iPGM, the reaction shown in Figure 2A, is further detailed.

**Figure 1.**
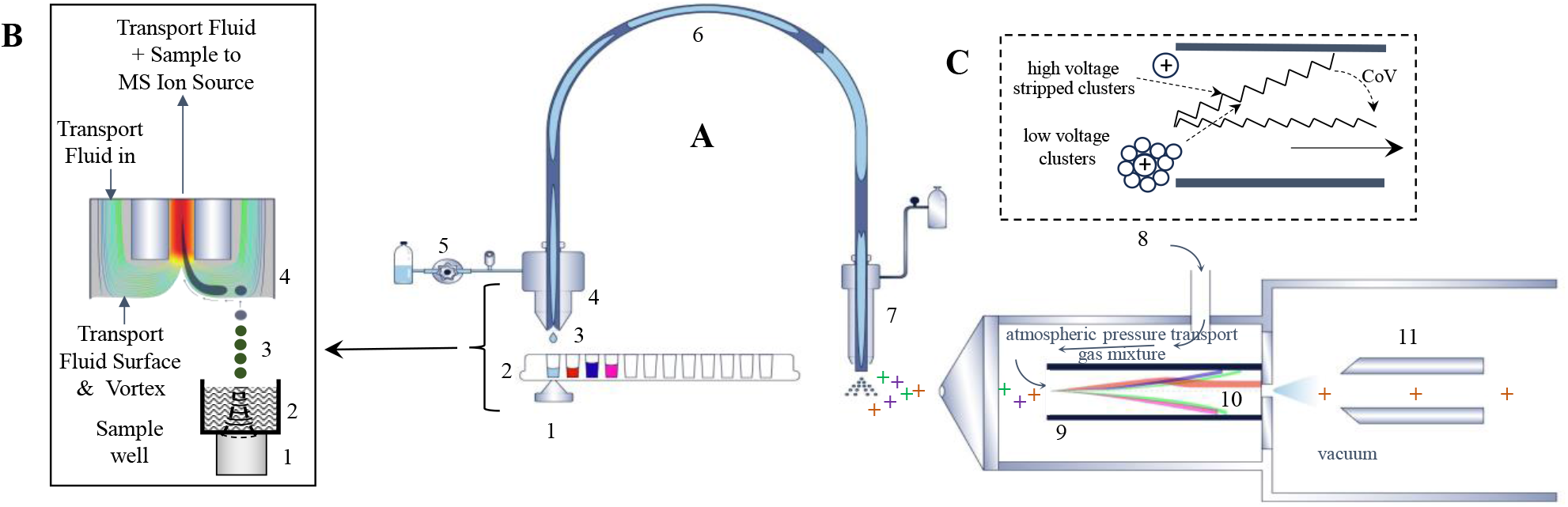
**(A)** DAEMS system. (1) Acoustic transducer dispensing sample droplets. (2) 1536-well microtiter plate (2-5uL/well) on x-y stage. (3) 2.5 nL sample droplets. One drop/sample for HTS at 1-2 sec/sample. (4) OPI captures, dilutes, and transports sample droplet. (**B**) Expansion of 1-4 in (**A**). (5) Transport fluid pump. (6) Diluted sample plug in transport fluid. (7) Pneumatically assisted electrospray ion source. Nebulizer gas expansion aspirates transport fluid from the OPI. (8) Vaporized solvents added to the nitrogen transport gas. (9) Planar differential ion mobility cell at atmospheric pressure in ion source, transport gas drawn by MS vacuum. RF voltage having a high and low voltage component, of different magnitude and duration, on each period of the sinusoidal waveform applied to planar electrodes. (10) Trajectory of 5 ions having dissimilar differential ion mobilities. **(C)** Expansion of 9 & 10. Sawtooth trajectory of ions and their clusters in Rf field migrate toward walls and discharge unless a DC compensation voltage (CoV) is used to steer on axis. Magnitude of CoV depends on rate ions and clusters migrate off axis^24^. Clustering during low voltage and de-clustering during high voltage period of waveform amplifies the differential ion mobility and accelerates ions to electrode. (11) Ion entrance optics of MS.

**Figure 2.**
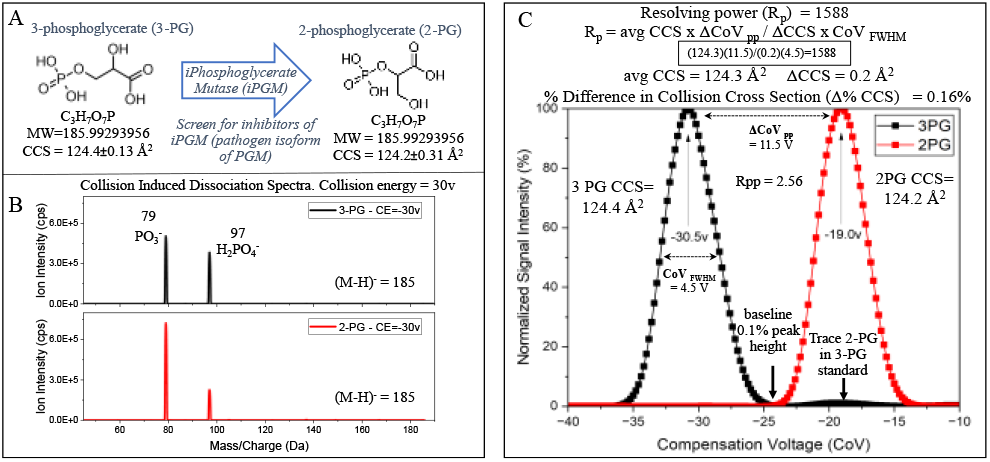
**(A)** The phosphoglycerate mutase reaction for HTS screening. **(B)** The CID spectra of 2 & 3-PG showing no distinguishable features. **(C)** Superimposed ionograms of 2 & 3-PG reference standards using 1% isopropanol in N_2_ acquired separately. Separation field 122 Td, MRM transition m/z 185 → 79. Variables in R_p_ calculation detailed. Ionogram acquired by acoustic infusion requiring successive 2.5 nL droplets at 10 Hz producing a continuous flow of analyte. Less than 1% 2-PG is observed in 3-PG standard commensurate with supplier specifications. CoV alternates between -30.5 & -19.0V during assay.

Internal standards were not used for quantification to avoid interference with the enzymatic reaction. The stability of the AEMS ion current, the volumetric precision of the droplet ejections, and elimination of sample preparation make this a unique feature among current HTMS technologies. Accurate mass measurements of the molecular ions or collision induced dissociation (CID) fragments cannot distinguish the isomers (Figure 2B) requiring separation prior to MS. DMS was chosen because its speed is compatible with HTMS sample introduction rates, and it has demonstrated the potential to separate structural isomers of relatively low molecular weight typically found in the metabolome.

AEMS employs MHz frequency sound waves focused through the bottom of microtiter plate wells to the sample surface transferring energy to the meniscus, disrupting surface tension, and frequencies and power were adjusted to provide a 2.5 nL sample droplet at a maximum of 1 second intervals to achieve the desired throughput. Speeds of 6 samples/s have been achieved for HTS assay’s^17^ requiring modifications of the OPI to lower flow resistance and enable higher transport fluid flow rates. The water-soluble PG analytes require a high aqueous transport fluid during droplet capture, dilution (∼1000x), and transport to a pneumatically assisted electrospray ion source as depicted in Figure 1A&B. The higher viscosity aqueous fluid requires a transfer tube/electrode with lower flow resistance to maintain a high fluid flow rate. The dilution reduces the ionization suppression from biological samples that are not purified before introduction into the MS.

The samples produce Gaussian shaped peaks of 1 second width at baseline as shown in subsequent Figures. For precise peak area integration, 8 data points are required for each compound particularly if external calibration is used^31, 39^. The number of different ions that can be monitored in a single sample ejection is determined by the DMS flight time and associated overheads while maintaining the required 8 measurements over these 1s peaks.

The 3 cm DMS cell requires ∼4.5 ms to fill as determined by the transport gas velocity drawn through by the MS vacuum. An additional 5 ms is required to fill the ion entrance optics of the MS and 2.5 ms for MRM detection (dwell) and voltage stabilization resulting in a minimum data acquisition time of ∼12 ms per analyte. This means that 10 analytes having the same MRM transitions can be monitored in each sample while acquiring the 8 data points across a 1 s peak. In this study a 50 ms dwell time with a pause time of 15ms was used because only 2 isomers were monitored providing ca. eight data acquisition points per peak for each isomer.

Low field mobility has a multiplexing advantage as all ions that do not diffuse to the walls are transmitted and detected from a single pulse of ions when synchronized to a TOF analyzer. To profile the 1-s wide peaks a maximum low field mobility flight time of 100 ms is required during which the resolution needed for the separation needs to be achieved. To date the required resolution at this speed has not been reported with low field mobility, particularly for launching a sample containing droplet of reproducible volume in the low nL range as illustrated in Figure 1 A&B^38^. In this study sound wave low mass isomeric pairs commonly encountered in the metabolome but can be achieved with DMS.

### Differential Ion Mobility

*Resolving Power and Mechanism of Separation*. The specificity of DMS separations is primarily derived from the thermodynamics of the clustering reactions and not on differences in the CCS of ions. Isomeric pairs in the metabolome have sufficiently different chemical properties to cluster with different numbers and bond strengths to neutral polar molecules ^25^. Since gas phase reactions are both specific and fast, high-resolution separations can be achieved in short periods of time.

Figure 2C shows the separation of 2 & 3-PG using a N_2_ transport gas containing IPA (Supplemental Info 4). Using the average single field CCS’s reported by Nicoles^1^ (Figure 2A), a calculated resolving power (R_p_) of 1588 is obtained with details in Figure.

Chorismate mutase and triosephosphate isomerase are potential drug targets, the separation of their substrates and products shown in Figure 3 A&B using ethyl acetate and isopropanol respectively. Since no literature CCS values are available the Rp is not calculated. In Figure 3 A&B and 2C a small signal (1-10%) from each standard appears at the CoV of its isomeric pair. This is because the reference standards used contain a small percentage of the corresponding isomer. More purified forms of these chemicals will improve the assay dynamic range.

**Figure 3.**
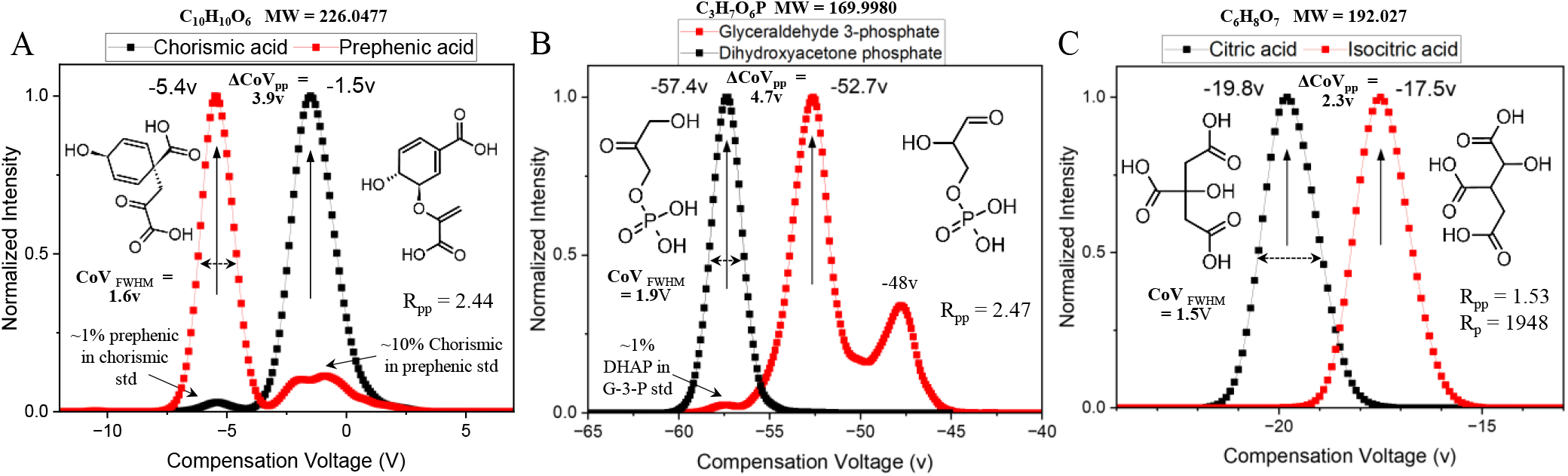
Superimposed ionograms of **(A)** chorismate mutase reaction isomers using 3% ethyl acetate and separation field 105 Td. **(B)** triosephosphate isomerase reaction isomers using 3% isopropanol and separation field 113 Td. **(C)** aconitase isomerase reaction isomers using 3% isopropanol and separation field 139 Td. Resolving gas of 1.95 and 1.44 slpm used for A&C and B, respectively. Two-peak resolutions, R_pp_, listed for each pair. Rp indicated only for the citrate/isocitrate separation. Reference standards for A and B contain trace amounts of the other isomer.

An additional peak appears in the G-3-P ionogram of Figure 3B at a CoV of -48 V. Different charge locations on the molecule will influence clustering as has been reported for prototropic isomers^25, 40-43^. Protonation of an atom that induces chirality can also lead to multiple peaks in ionograms where diastereomers are created and exhibit unique cluster conformations resolvable by differential ion mobility^25^. It is hypothesized that this additional peak in Figure 3B is the negative ion equivalent of a protomer having the negative charge on a different phosphate oxygen. The elucidation of the identity of this peak is being pursued employing HPLC separations prior to DMS and accurate mass TOF CID and EAD spectra to probe its structure. Nevertheless, the presence of this peak will not interfere with quantitative measurements of the main components.

The substrates and products of the aconitase reaction, citrate and isocitrate, ave reported CCS’s t at differ by 0 08 ^1^. The calculate R_p_ is 1948 shown in Figure 3C. To our knowledge this is the highest mobility resolving power reported to date for compounds from the metabolome.

A two-peak resolution value can be useful for assessing if both isomers can be quantitated in the presence of high concentrations of the other. Adapting the equation of Tabrizchi^44^, 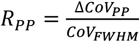, the two-peak resolution for the data in Figure 3A,B, & C are 2.44, 2.47, 1.53. respectively and 2.56 for the data in Figure 2C. Inspection of the data indicates that the isomers can be independently monitored at these two-peak resolution values accounting for the fact that the reference standards used for 3-PG, chorismic acid, prephenic acid, and glyceraldehyde 3-phosphate contain a small percentage of the other isomer.

Literature CCS values can vary depending on the experimental conditions. Nicholes^1^ and Zheng^32^ reported CCS’s for 2 which would lead to lower calculated R_p_ values than those presented here. Resolving power based on CCS differences is a meaningful and useful mobility metric for comparing different instrument configurations and for considering the utility of CCS databases to aid in the identification of unknowns. The two peak method provides some indication if separation of targeted analytes for quantitation purposes is possible. It is interesting to note that among the 4 examples shown here the citrate/isocitrate isomeric pair produced the highest resolving power in CCS space and the lowest using the 2 peak resolution calculation suggesting that the most appropriate resolution calculation to use depends on the application to which it is referenced as alluded to by the Bard.

“And thus t e native hue of resolution

Is sicklied o’er with the pale cast of thought

“Shakespeare. Hamlet, Scene 1, Act 3.

The high resolution is due to the differences in the gas phase clustering properties of the isomers. As shown in Figure 1C, during the high voltage period of the waveform the ion is stripped of its neutral solvent cluster molecules by the high energy collisions thereby increasing its velocity toward the wall. During the low voltage component of the waveform period, the ion and neutral solvent molecules re-cluster, reducing t e aggregate’s velocity in t e other direction. The number of neutral molecules to cluster and their bond strength depends on the chemical properties of the ion. The repetitive clustering and declustering occurs ∼10,000 times during the < 5 msec transit through the cell at atmospheric pressure. It is the difference in these two velocities that determine the net velocity toward the electrode which in turn determines the CoV required to steer an ion back on axis, thus the name differential ion mobility. A highly clustered ion with strong bond strengths will require a greater CoV to correct its trajectory^25^.

Polar low molecular weight molecules, such as those commonly found in the metabolome, readily form ion-molecule clusters with polar solvents. This explains the high resolving powers observed for components of the metabolome and other polar low molecular weight compounds where a few solvent clusters have a large relative effect on its differential mobility (Supplemental Info 2 comparing with/wo modifier).

The chemical specificity of the clustering process is illustrated in Figure 3 where different vapors are used to induce separations (Supplemental 3 & 4 for modifier effect on 2-& 3-PG separation). The situation is somewhat analogous to HPLC where chemistry is optimized to effect separations. HPLC has multiple phases (liquid and stationary) and chemistries with which to leverage chemical differences for separation purposes giving it a significant resolution and peak capacity advantage over the gas phase processes occurring in DMS. But HPLC cannot match the speed of DMS, a critical consideration for separation tools being used to hyphenate with emerging HTMS sample introduction technologies.

An additional benefit the separation capability of DMS provides HTMS is ion filtration prior to the MS. DMS filtering has been shown to reduce MS analyzer contamination by 10-fold because greater than 95% of the background ion current doesn’t enter the ion optics in vacuum^45^. This feature is particularly valuable for instruments running several tens of thousands of samples per day for weeks as is the case with HTS primary library screens.

### iPGM Assay Analytical Metrics

*LOQ, Dynamic Range, Precision, Accuracy, and HT Quant*. The substrates and products of the reaction shown in Figure 2A are being simultaneously monitored. Figure 4A shows a calibration ladder and Figure 4B a calibration curve for 3-PG in the assay buffer across a concentration range typically used for the bioluminescent assay. The reaction conditions and buffers were the same for DAEMS and the bioluminescent assay. The composition of the reaction ejected into the MS is listed in Figure 4C providing a suppression signal loss of ∼2x compared to water standards. This illustrates that quantitation without sample preparation is feasible for this assay. The dilution of ∼1000-fold that occurs in the OPI minimizes ionization suppression making sample preparation for matrices with these salt levels unnecessary^36, 46^. The limit of detection estimate from this data is 50 nM however avenues for improving that by an order of magnitude exist outside the scope of this report. This would involve using mass spectrometers that can accommodate higher ion and gas loads^5^.

**Figure 4.**
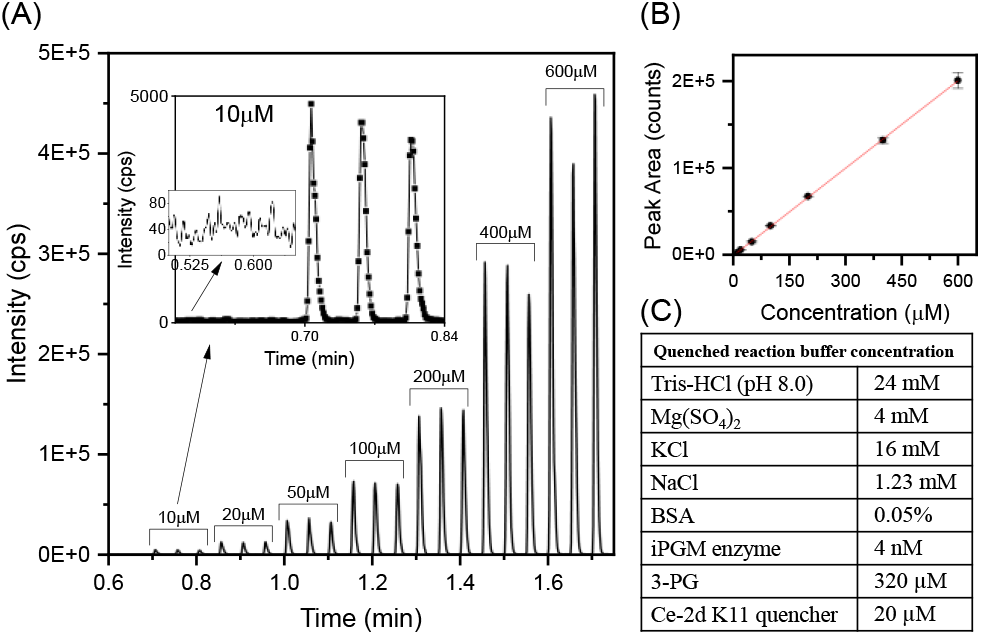
(A) Chronogram of 3-PG calibration curve covering the assay dynamic range. CoV switching between -19.5 & -30V and MRM (M-H)^-^ 186→ 79. Triplicate 2.5nL injections per concentration with expansion around the lowest calibration point and the background level. (B) Calibration curve based on peak areas (no internal standard) from the data in (A). (C) The DAEMS assay buffer composition.

AEMS has demonstrated the ability to maintain consistent peak areas for tens of thousands of samples^47^. This presents the possibility that internal standards will not be required for concentration determinations, an important consideration with HTS screens and other applications where internal standards can interfere with the reactions or be otherwise impractical. Figure 5 demonstrates a peak area CV of 8.3% over the course of a 1536 well plate. Since there is no sample preparation involved any additional variability would be limited to reagent dispensing errors. This allows for an external calibration approach referred to as HT Quant.

**Figure 5.**
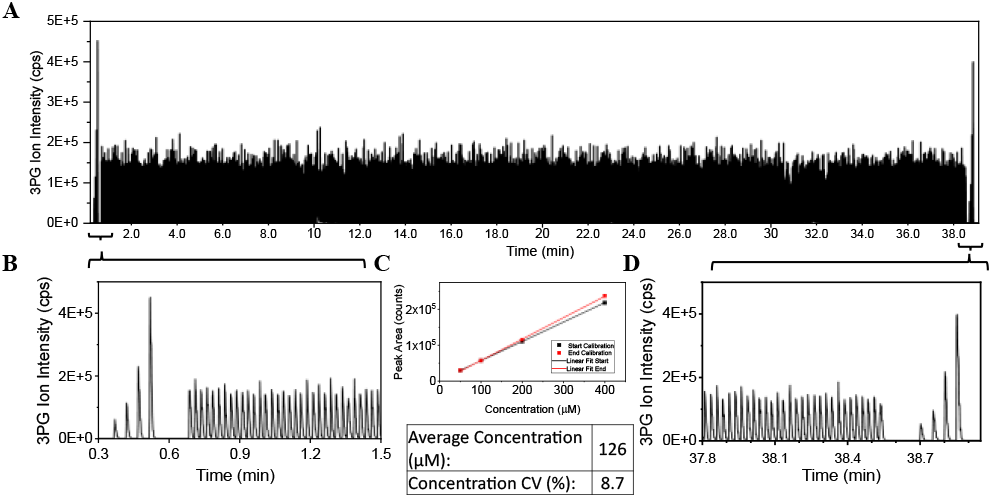
HT Quant. External standard quantitation using calibration curves at the beginning and end of a 1536 well plate run. (**A)** 1536-well plate at 1 sample/1.5 sec of a 3-PG QC in the assay buffer. sandwiched between calibration sample runs. (**B&D)**. Expansion of calibration curve samples at beginning and end of the plate. (C) Overlay of the two calibration curves and QC concentration statistics.

Two calibration curves from the first and last four wells straddle the plate to account for possible signal drift over the course of the run as shown in Figure 5A. It has been demonstrated with other HTS assays using AEMS that signal stability can be maintained over the course of ∼20 plates (∼ 30,000 samples) and can be readily corrected with a quick water wash of the OPI capture port^13^. Figure 5B & D expands the x-axis of the calibration regions. Figure 5C shows that the two external calibration curves remain correlated during this 1536 well plate run. Each plate would repeat the two calibration curves requiring a minimal amount of time/plate, 8 calibrants in 8 seconds at 1 Hz. The noticeable pattern of these calibration curves also establishes whether the first and last wells were successfully dispensed. This avoids any possibility of a frame shift error if, for example, the first well was not dispensed. This approach has been successful for quantitation of pharmaceutical library components without the use of an internal standard^48^.

### Comparison of DAEMS to Bioluminescence iPGM Assay

*Z’ and IC50*. T e Z’s for t e biolu inescence coupled-enzyme and DAEMS assays run at four different incubation times are provided in Figure 6A & B and are shown to be similar and of high quality (Supplemental Info 5 for detail).

**Figure 6.**
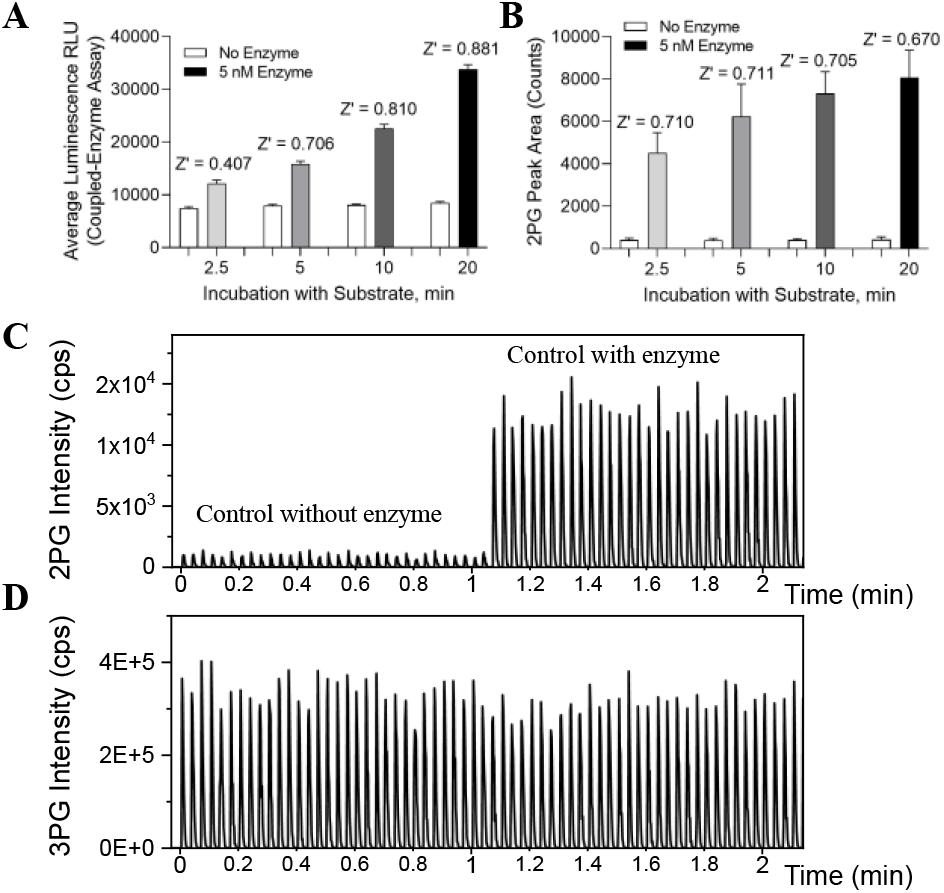
Data from controls with enzyme to catalyze conversion and without enzyme for no conversion. **(A)** Average intensity plots from the biolu inescence assay used to calculate Z’ **(B)** Average intensity plots fro t e DAEMS assay used to calculate Z’ Error bars represent the standard deviation of 32 replicate wells. **(C)** DAEMS generated chronogram of 2-PG from the two controls used to calculate Z’ with reaction incubation time of 2.5 minutes. **(D)** Chronogram of the 3-PG signal acquired at same time as in (C).

The short 2.5-minute incubation time produced a low signal in the bioluminescence assay resulting in a Z’ outside t e acceptable range necessitating longer incubations. The high bioluminescence background, as well as the lowered signal, contributes to the S/N reduction. The higher DAEMS sensitivity allows shorter incubation times thus higher throughputs.

The chronogram of the controls in Figure 6C establishes the noise level used to determine the S/N for the Z’ calculation. The 2-PG background peaks in the control (0-1.05 min) are due to a ∼0.25% contamination of the 3-PG standard with 2-PG also estimated by the chemical manufacturer. A more purified 3-PG standard without 2-PG would increase the DAEMS S/N by 10x further i proving t e assay Z’ and dyna ic range The control with enzyme (1.2 to 2.1 min) shows a 5% conversion of 3-PG to 2-PG for the 2.5-minute reaction time as expected.

The bioluminescent assay was previously used to determine the IC_50_ values of a class of cyclic peptides known as ipglycermides which were discovered by iPGM affinity selection from a vast RNA-encoded cyclic peptide library^33^. The DAEMS titration dose response curves used to determine the IC_50_’s for t ree of t ese in hibitors are shown in Figure 7A-C.

**Figure 7.**
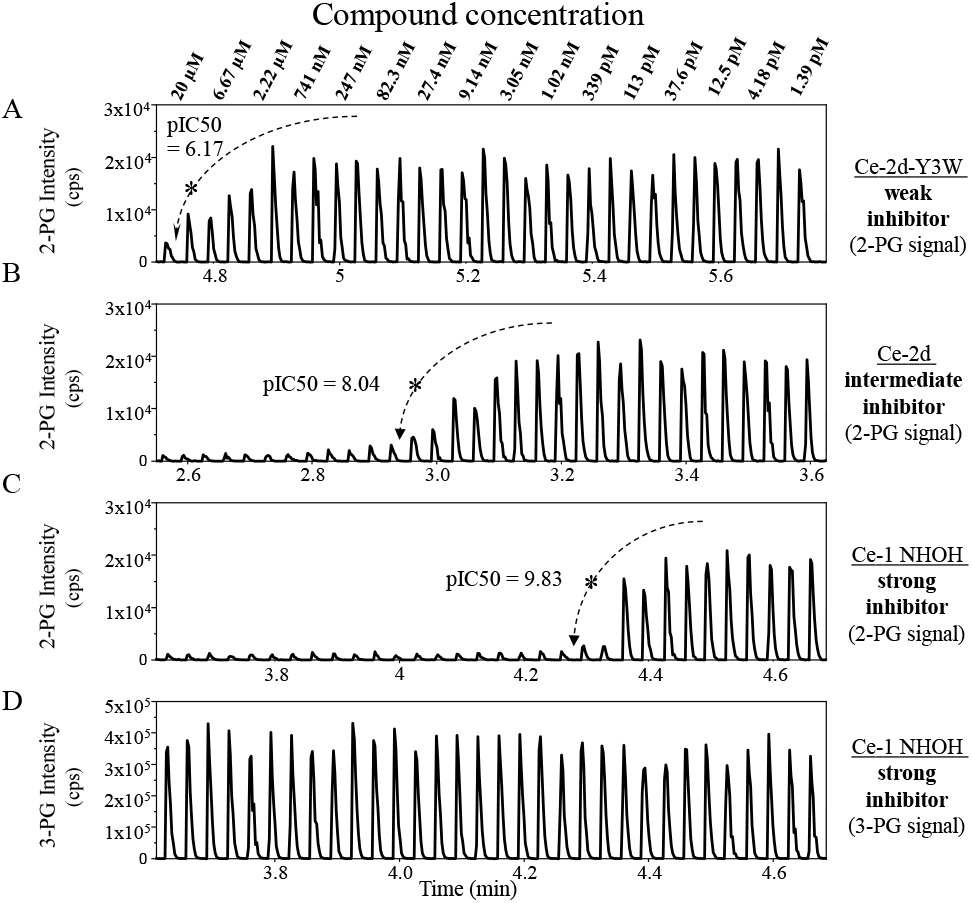
**A, B, C**. Extracted ion current chronograms (XIC) for 2-PG as the concentration of 3 different inhibitors is titrated from 20 µM to 1.39 pM to deter ine IC50’s **A**. Ce-2d-Y3W weak inhibitor. **B**. Ce-2d intermediate inhibitor. **C**. Ce-1 NHOH strong inhibitor. **D**. The XIC for the 3-PG monitored during C. Reactions quenched at a 5-min.

The chronograms in 7 A-C show a clear trend toward increasing inhibition at lower concentrations with stronger inhibitors. The 2-PG signal change is relatively large going from baseline, as limited by the background 2-PG in the 3-PG standard (750 nM), to 5% conversion (15 µM). It should also be noted that since only ∼ 5% of the 3-PG is consumed the change in substrate concentration is not visually obvious but measurable based on peak areas (Figure 7D).

The graphs in Figure 8A and B show the inhibitor concentration response curves for DAEMS and the bioluminescent assays determined at the 4 incubation times. There is no significant difference in the pIC_50_ values although the DAEMS assay shows a slight shift toward lower pIC_50_ at shorter reaction times (Fig 8A). The luminescence assay shows consistent measurements across reaction times (Fig 8B). The reason for the increase in IC_50_’s it longer incubation times for DAEMS is likely because the product of the reaction (2-PG) accumulates which shifts the reaction equilibrium. The 2-PG product is consumed in the bioluminescence assay increasing the duration of the linear kinetic range of this reaction.

**Figure 8.**
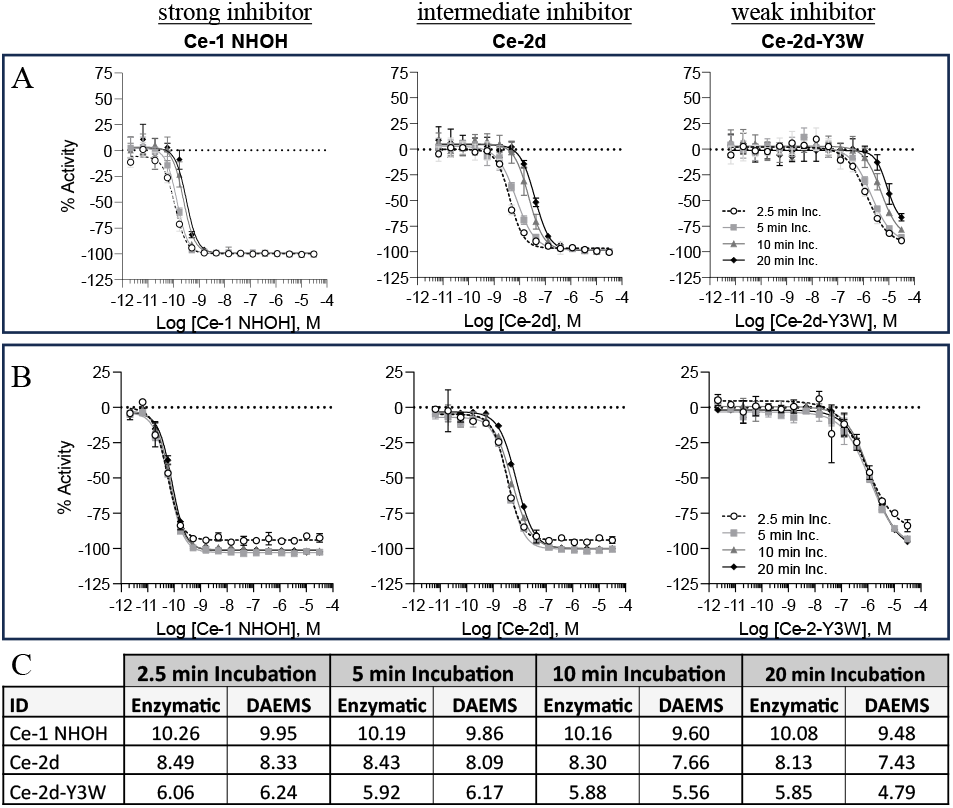
Concentration response curves (CRCs) for three iPGM inhibitors measured at four incubation times fit to the Hill equation (Supplemental Info 6&7). **(A)** DAEMS assay showing standard mean error of six replicates. **(B)** Bioluminescence assay. Standard mean error for two replicates per concentration. **(C)** Comparison of pIC_50_.

### Real Time Enzyme Reaction Monitoring

The possibility to use DAEMS to probe the kinetics of the iPGM reaction was measured and compared to a UV absorbance-based coupled-enzyme assay. Since the bioluminescence assay only provides reaction end-point data, an absorbance-based NADPH depletion assay was developed and used for kinetic studies as previously reported^33^. DAEMS can be used for both endpoint and kinetic measurements.

Since the sampling volume is low (2.5 nL) and no intermediate sample transfer, processing, or plating is required, the progress of the reaction can be monitored in real time at frequencies ranging from sub-second to hours^36^. The data in Figure 9 is a 5 second sampling interval. The depletion of NADPH in the absorbance-based assay requires upwards of 10 minutes to complete the linear phase of the coupled-enzyme reaction. DAEMS detects 2-PG within the first minute and reaches equilibrium in less than 5 minutes.

**Figure 9.**
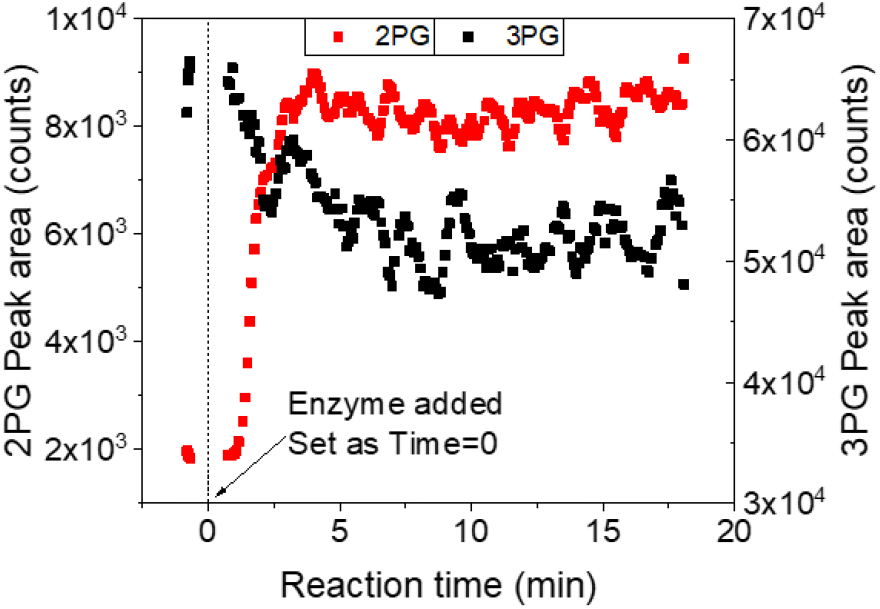
Real-time reaction monitoring of the production of 2-PG and the consumption of 3-PG to establish iPGM reaction kinetics. 2.5 nL dispensed every 5 sec for 20 minutes (240 measurements), consuming 0.6 µL. 5 nM *C. elegans* iPGM enzyme and 400 µM 3-PG substrate in buffer at ambient temperature. 5-point adjacent-averaging smooth.

A continuous stream of the reaction with no time gaps can be generated with multiple 2.5 nL droplets ejected at 20 Hz with a sample consumption rate of 50nL/s (3 µL/minute). This provides the opportunity to probe mechanistic details of enzymatic reactions identifying and establishing the kinetics of transient intermediates which was earlier shown to be feasible with syringe infusion-based techniques^49-50^ and is only possible using real-time reaction monitoring and mass spectrometric detection.

## CONCLUSIONS

We report the first implementation of ion mobility mass spectrometry with an ultra-high throughput sample introduction technology for HTS of isomers typically encountered in the metabolome. The system involves integrating DMS with AEMS (DAEMS). It provides the capability to simultaneously quantitate the concentration of structural isomers that have identical masses and fragmentation spectra at sample introduction speeds of ≥ 1 second per sample. Its potential as a label free approach to high throughput screening of metabolic enzymes whose substrates and products are isomers, such as isomerases, mutases, and transferases, is de onstrated by co paring t e Z’ and IC_50_ determinations from DAEMS to a well-established coupled enzyme bioluminescent assay for the isoform of phosphoglycerate mutase that is a target for a variety of microbial and parasite borne diseases. The assay metrics are shown to be comparable along with demonstrating many of the advantages of a label free approach over the coupled-enzyme bioluminescence assay.

A large percentage of the metabolome is composed of isomeric combinations as indicated by the composition of the MSMLS library wherein 30% of its components are isomeric species^1^. The potential of DAEMS as a general approach for screening is indicated by showing the separation of two additional promising targets, chorismate mutase and triosephosphate isomerase. Resolving powers as measured in CCS space for 2 & 3-phosphoglycerate and the citrate – isocitrate isomeric pair were calculated from the ionograms and published drift tube CCS values to be 1588 and 1948 respectively, the highest values reported in the mobility literature for compounds found in the metabolome. These high resolutions provide sufficient separation for quantitation over a broad dynamic range without cross channel overlap of signal.

For HTS, particularly as it relates to quantification of low molecular weight isomeric species typical of the metabolome, mobility resolving po ers > 00 need to be ac ieved in the tens of millisecond range. To date this has not been demonstrated with low field IM-MS but demonstrated here using DMS.

AEMS based label free HTS assays have been used for an increasing number of primary screens of pharmaceutical libraries because of the advantages over parallel photometric detection systems. A reduction in false positives and negatives occurs with direct measurement because common interferences such as compound autofluorescence, light scattering, fluorescence quenching, and inhibition effects of the coupled enzymes are eliminated. In addition, lower limits of quantitation, a broader dynamic range, low sample consumption, reduced time and complexity for assay development, and lower consumable reagent costs are achieved.

This study represents the first step in validating DAEMS for primary library screens where isomer separation is required. Future experiments will focus on full library screens for inhibitors of promising isomerase targets as those described here in. Additionally, DMS has demonstrated the ability to minimize mass spectrometer ion optics contamination under severe sample loads particularly relevant to large scale library HTS screening campaigns. It is anticipated that DAEMS will represent a significant advancement in the general field of high throughput mass spectrometry addressing the challenge of simultaneously quantifying at high-speed structural isomers and other isobaric interferences as well as dealing with instrument contamination issues that are inevitable with systems analyzing upwards of 100,000 samples per day for extended time periods.

## ASSOCIATED CONTENT

### Supporting Information

Impact of OPI transfer fluid composition on AEMS peaks (SI 1); DMS separation of 2-, 3-PG without modifier (SI 2); DMS separation of 2-, 3-PG using Methanol as modifier (SI 3); DMS separation of 2-, 3-PG at different concentrations of Isopropanol as modifier (SI 4); Detailed statistics for luminescence and DAEMS assays (SI 5); Concentration response curves for all the runs individually (SI 6); Calculated parameters from the concentration response curves (SI 7); Protocol that was followed for coupled-enzyme and DAMES assays (SI 8); Compound plate layout and plate images (SI 9); Reagents and materials used (SI 10).

## AUTHOR INFORMATION

### Notes

DAEMS is not a commercial product. SCIEX is the commercial provider of Echo MS, Echo MS+, SelexION, SelexION+, and the 6500+ QTRAP mass spectrometer.

## ACKNOWLEDGMENT

This work was done under the NIH/NCATS – SCIEX CRADA #2020-08f under auspices of Sam Michael, Chief, Information Technology Resources Branch (ITRB). This research was supported in part by the Intramural Research Programs of the National Center for Advancing Translational Sciences, NIH under project 1ZIATR000053 (J.I.). The content of this publication does not necessarily reflect the views or policies of the Department of Health and Human Services, nor does mention of trade names, commercial products, or organizations imply endorsement by the U.S. Government.

We thank Jay Corr and Yves Le Blanc for their support and the fruitful discussions, Stanislaw Potyrala for building OPI electrodes, and Lyle Burton for help with the data processing software.

