## Supplementary Information for "A Differential Ion Mobility Acoustic Ejection Mass Spectrometer System for Screening Isomerization-Mediating Enzyme Drug Targets"

A

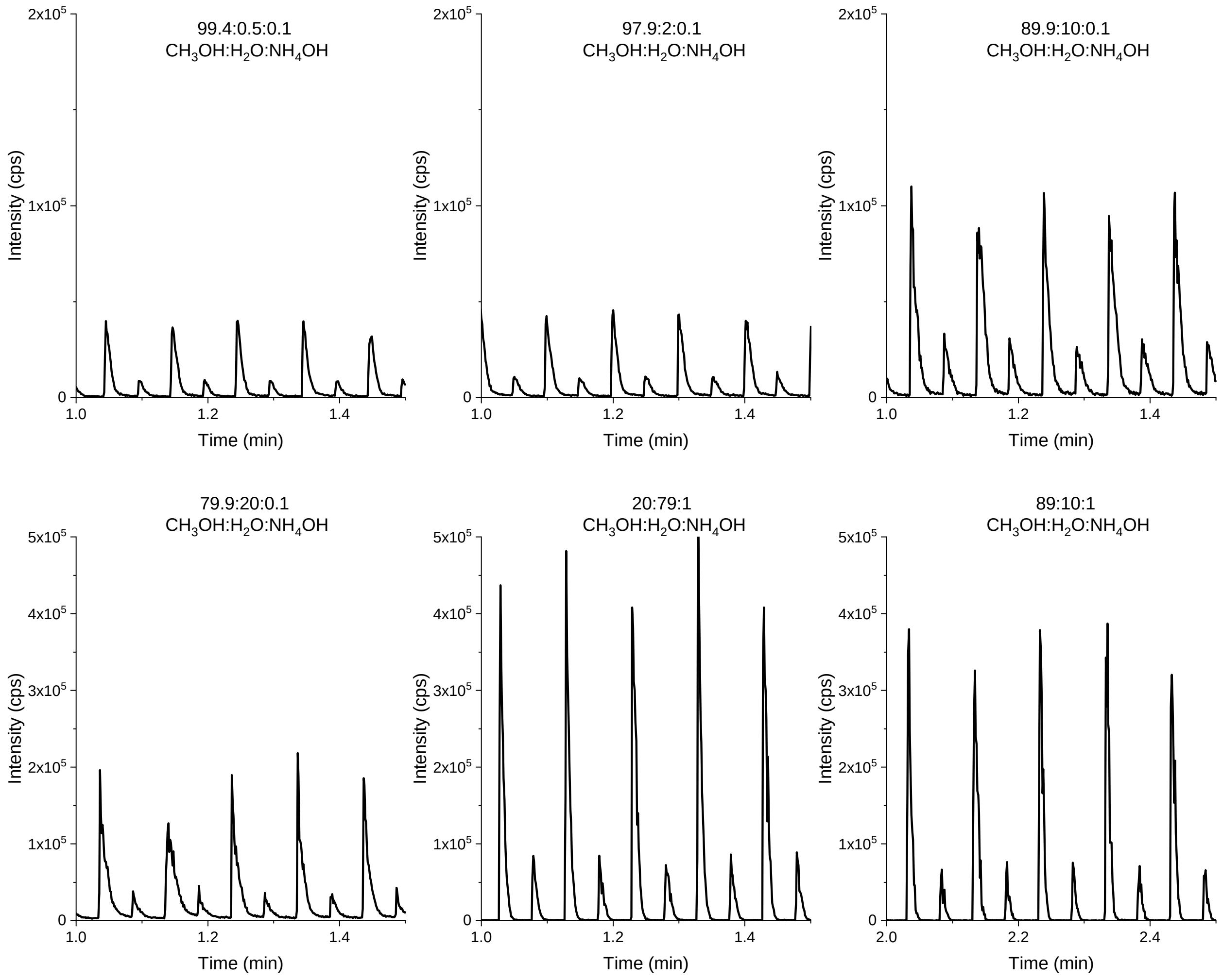


B

To reduce the adhesion of these polar compounds to the glass surface of the PEEKsil tubing for the OPI transfer line, a PEEK transfer line was used instead with the same nominal internal diameter as the PEEKsil tubing. To reduce the flow resistance of the tubing for the high aqueous transport fluid as shown in the figure, a shorter emitter of ca. 24mm was used instead of the standard 80mm long emitter. This short emitter was press-fit into the PEEK tubing by about 3mm.

**Supplementary info** 1- (A) Impact of OPI transfer fluid composition on analyte solubility and consequently peak shape, width, and intensity. Samples of 100 µM and 400 µM of 3-PG in the assay buffer were diluted 2x in deionized water and alternating 2.5 nL droplets were ejected from the samples every 3s. Y-axes were scaled differently between the top and bottom plots for optimal visualization of the data. (B) Description of the changes made to the ESI emitter/transport tubing.

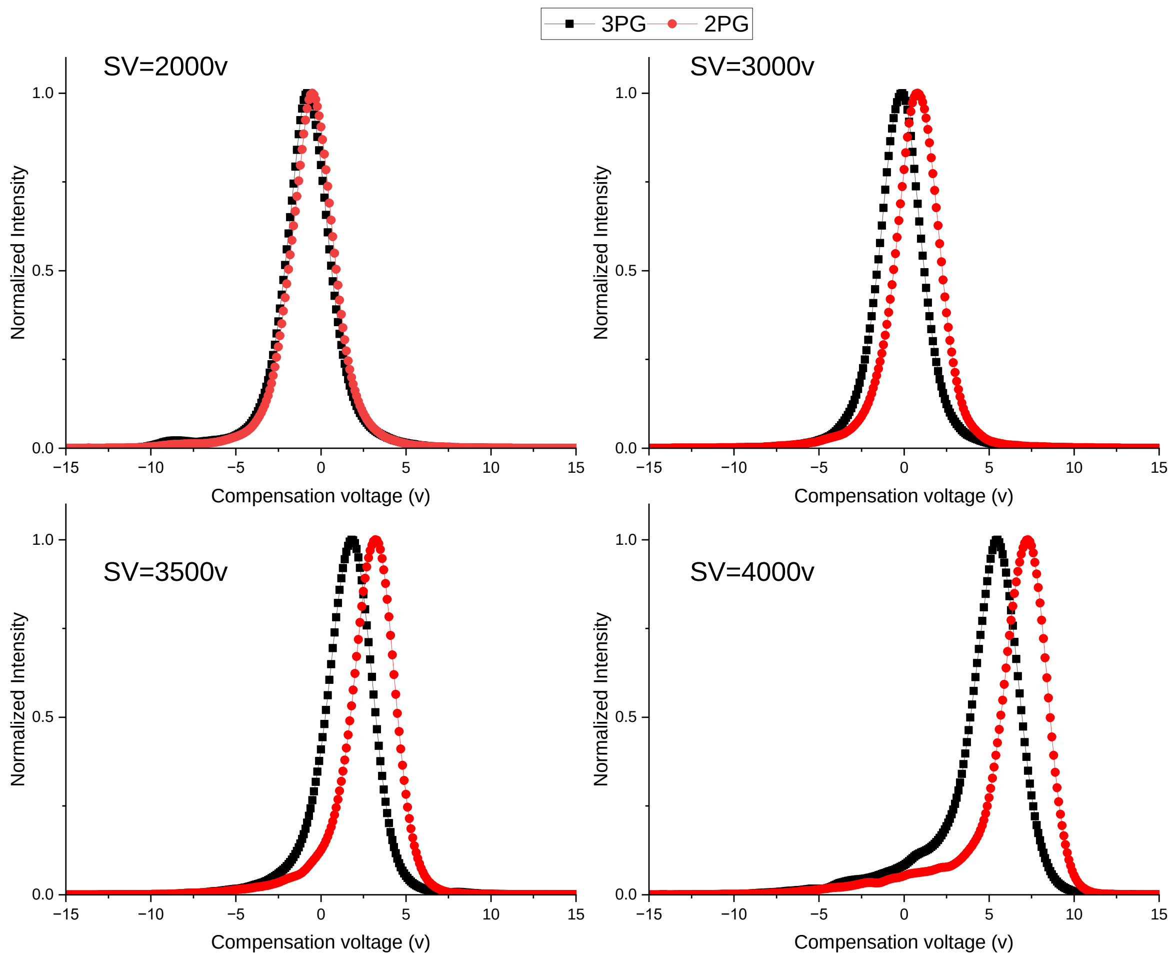


**Supplementary info 2-**Compensation voltage ramps at different SV values without the use of a modifier indicating an insufficient peak separation. Samples were infused by syringe.

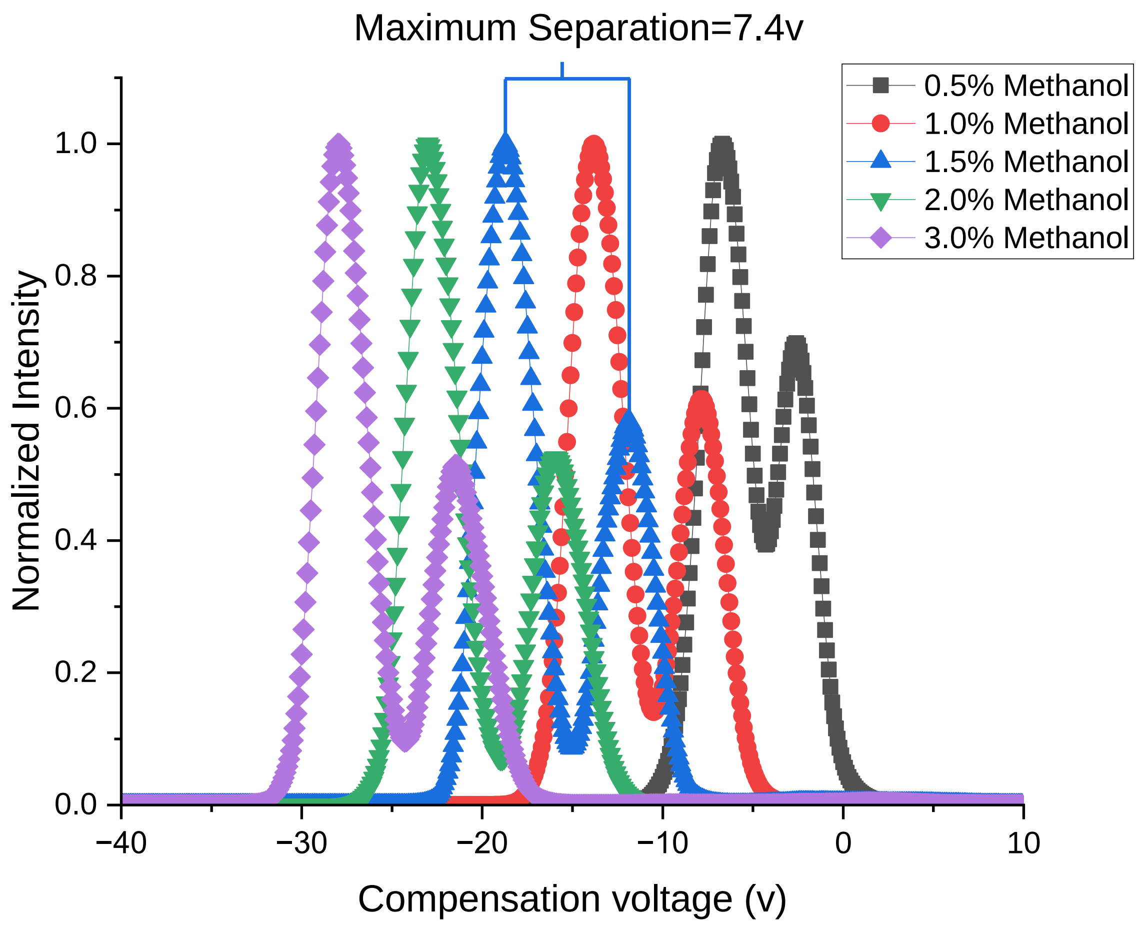


**Supplementary info 3-**Compensation voltage ramps at different concentrations of methanol as a modifier for a mixture of 2-PG and 3-PG. Max peak separation of 7.4v was achieved for 1.5% Methanol. Samples were run with acoustic droplet ejection at a rate of 10 Hz for pseudo-infusion mode and Separating Voltage=3500v. Left-side peak in each trace is 3-PG and the right-side peak is 2-PG.

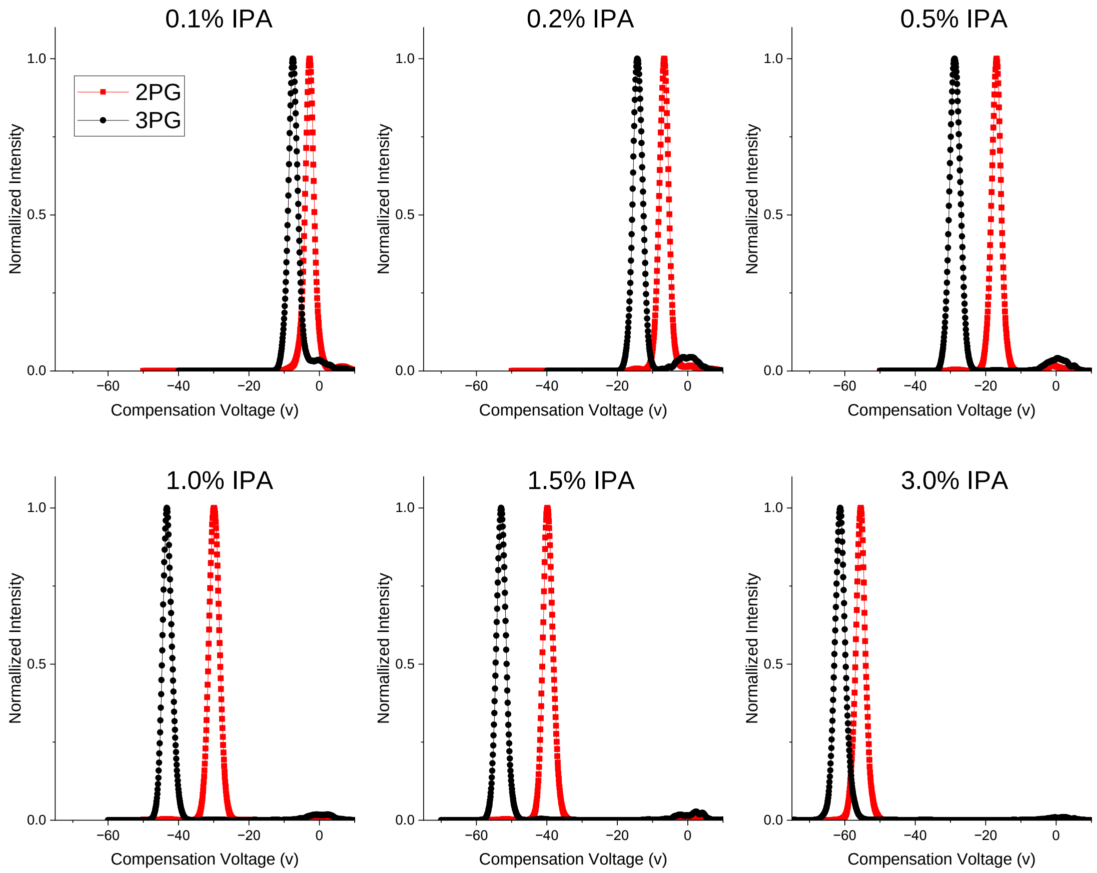


**Supplementary info 4-**Compensation voltage ramps at different concentrations of Isopropanol (IPA) as a modifier. 3-PG and 2-PG peaks are baseline resolved with the maximum peak separation achieved at 1.0% IPA. 2-PG and 3-PG Samples were prepared in DIW and were infused with a syringe,  Separating Voltage=3500v.

(A)

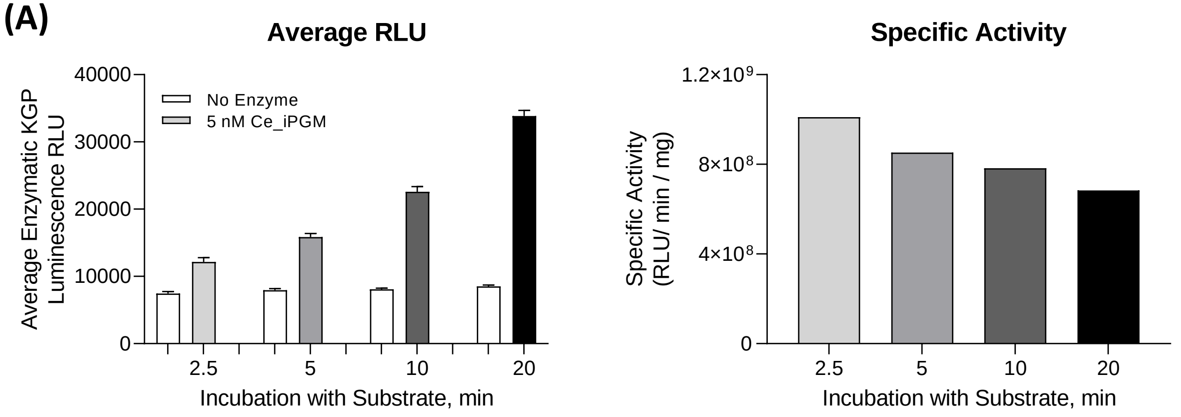


| | 2.5 min | | 5 min | | 10 min | | 20 min | |
|---|---|---|---|---|---|---|---|---|
| | No Enzyme | Enzyme + DMSO | No Enzyme | Enzyme + DMSO | No Enzyme | Enzyme + DMSO | No Enzyme | Enzyme + DMSO |
| Average | 7445.7 | 12138.3 | 7953.4 | 15855.8 | 8085.0 | 22589.9 | 8530.7 | 33847.9 |
| St. Dev. | 291.1 | 636.2 | 236.7 | 538.6 | 153.3 | 765.0 | 178.2 | 830.1 |
| %CV | 3.9 | 5.2 | 3.0 | 3.4 | 1.9 | 3.4 | 2.1 | 2.5 |
| S:B | | 1.6 | | 2.0 | | 2.8 | | 4.0 |
| S:N | | -6.7 | | -14.3 | | -19.4 | | -31.1 |
| Z' | | 0.407 | | 0.706 | | 0.810 | | 0.881 |

| *23 °C Ambient Temp* | 2.5 min | 5 min | 10 min | 20 min |
|---|---|---|---|---|
| Corrected Ave. RLU | 4692.6 | 7902.4 | 14504.9 | 25317.2 |
| Vol. added to 6ul Rx (ml) | 3.03E-06 | 3.03E-06 | 3.03E-06 | 3.03E-06 |
| Incubation Time (min) | 2.5 | 5 | 10 | 20 |
| Concentration (mg/ml) | 0.613 | 0.613 | 0.613 | 0.613 |
| Spec. Act. [(RLU/min/mg) | 1.01E+09 | 8.52E+08 | 7.82E+08 | 6.83E+08 |

(B)

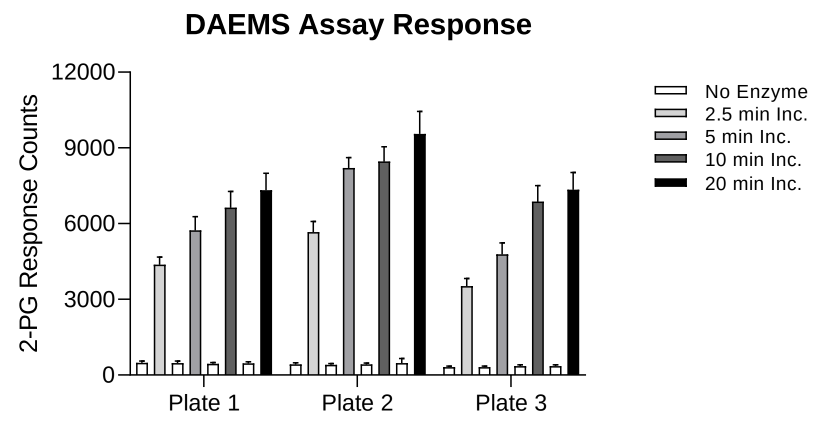


| | 2.5 min Incubation | | | | | | 5 min Incubation | | | | | |
|---|---|---|---|---|---|---|---|---|---|---|---|---|
| | Plate 1 | | Plate 2 | | Plate 3 | | Plate 1 | | Plate 2 | | Plate 3 | |
| | No Enzyme | Enzyme + DMSO | No Enzyme | Enzyme + DMSO | No Enzyme | Enzyme + DMSO | No Enzyme | Enzyme + DMSO | No Enzyme | Enzyme + DMSO | No Enzyme | Enzyme + DMSO |
| Average | 483.4 | 4373.2 | 429.3 | 5665.2 | 310.4 | 3528.3 | 478.6 | 5737.4 | 402.4 | 8201.3 | 311.0 | 4780.3 |
| St. Dev. | 68.3 | 299.7 | 57.3 | 417.6 | 40.9 | 297.2 | 73.9 | 536.5 | 57.4 | 416.9 | 43.1 | 458.7 |
| %CV | 14.1 | 6.9 | 13.4 | 7.4 | 13.2 | 8.4 | 15.4 | 9.4 | 14.3 | 5.1 | 13.8 | 9.6 |
| S:B | | 9.0 | | 13.2 | | 11.4 | | 12.0 | | 20.4 | | 15.4 |
| S:N | | -12.7 | | -12.4 | | -10.7 | | -9.7 | | -18.5 | | -9.7 |
| Z' | | 0.716 | | 0.728 | | 0.685 | | 0.652 | | 0.818 | | 0.663 |

| | 10 min Incubation | | | | | | 20 min Incubation | | | | | |
|---|---|---|---|---|---|---|---|---|---|---|---|---|
| | Plate 1 | | Plate 2 | | Plate 3 | | Plate 1 | | Plate 2 | | Plate 3 | |
| | No Enzyme | Enzyme + DMSO | No Enzyme | Enzyme + DMSO | No Enzyme | Enzyme + DMSO | No Enzyme | Enzyme + DMSO | No Enzyme | Enzyme + DMSO | No Enzyme | Enzyme + DMSO |
| Average | 440.0 | 6636.6 | 425.2 | 8461.5 | 352.0 | 6873.4 | 462.7 | 7320.7 | 470.8 | 9553.3 | 353.0 | 7345.3 |
| St. Dev. | 54.5 | 639.3 | 48.6 | 579.9 | 53.3 | 633.0 | 59.8 | 672.7 | 184.5 | 891.4 | 48.5 | 682.4 |
| %CV | 12.4 | 9.6 | 11.4 | 6.9 | 15.1 | 9.2 | 12.9 | 9.2 | 39.2 | 9.3 | 13.7 | 9.3 |
| S:B | | 15.1 | | 19.9 | | 19.5 | | 15.8 | | 20.3 | | 20.8 |
| S:N | | -9.7 | | -13.8 | | -10.3 | | -10.2 | | -10.0 | | -10.2 |
| Z' | | 0.664 | | 0.765 | | 0.684 | | 0.680 | | 0.645 | | 0.686 |

**Supplementary Info 5** - Assay statistics summary for *C. elegans* iPGM enzyme incubated with 3-PG substrate across a 20 min time course for **(A)** the coupled-enzyme functional luminescence assay measured with Kinase-Glo Plus (KGP) reagent and **(B)** the DAEMS assay. Control samples of 5 nM of enzyme or no enzyme buffer were treated with DMSO vehicle control and 20 mM Ce2d K11 peptide quencher added at designated time points. Average luminescence RLU (A) were compared to the 2-PG product response counts (B) between the assays. The specific activity of the enzyme across the time course is shown (A). Statistics were calculated for 32 replicates for each enzyme and no enzyme control per time point, per replicate plate in the DAEMS assay, and error bars represent standard deviation. The time points that produced statistically significant Z' >0.5 are highlighted in yellow.

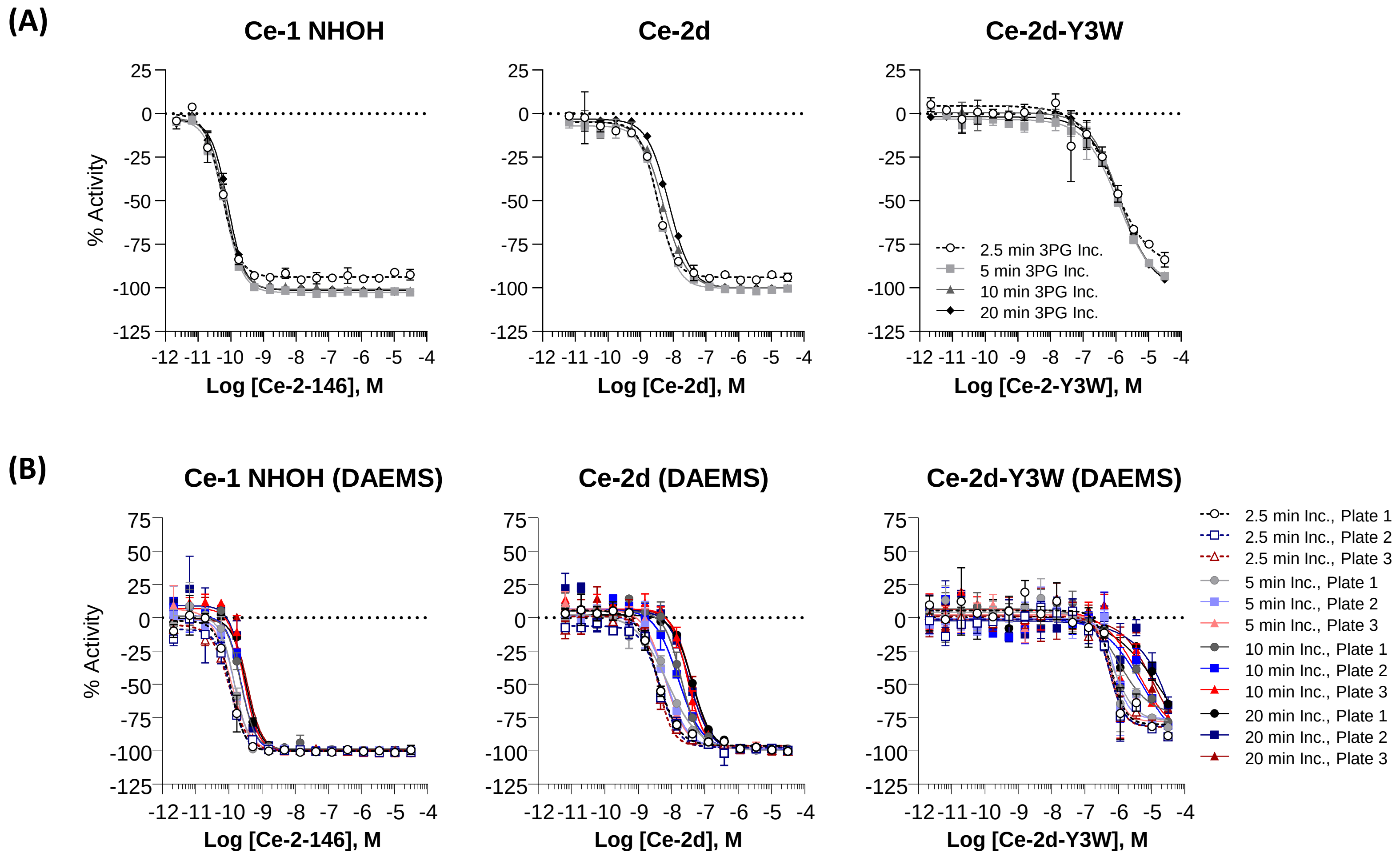


**Supplementary Info 6** - Concentration response curves (CRCs) for a 20 min time course tested across macrocyclic peptide inhibitors in **(A)** the coupled-enzyme Kinase-Glo Plus (KGP) luminescence end point assay and **(B)** the DAEMS assay. CRCs represent row-wise normalized data to DMSO neutral control and no enzyme control as -100% inhibition for each time point based on luminescence RLU for (A) and 2-PG response counts for (B). Curves were fit in GraphPad Prism and error bars represent standard deviation for two replicate wells per titration. The DAEMS Assay was run across three replicate plates.

**(A)**

| Sequence | ID | Enzymatic Assay, 2.5 min Max I | Enzymatic Assay, 2.5 min pIC50 | Enzymatic Assay, 5 min Max I | Enzymatic Assay, 5 min pIC50 | Enzymatic Assay, 10 min Max I | Enzymatic Assay, 10 min pIC50 | Enzymatic Assay, 20 min Max I | Enzymatic Assay, 20 min pIC50 |
|---|---|---|---|---|---|---|---|---|---|
| $^{D}$YDYPGDYCYLYGTC-NHOH | Ce-1 NHOH | -92.5 | 10.26 | -102.6 | 10.19 | -101.7 | 10.16 | -102.4 | 10.08 |
| $^{D}$YDYPGDYCYLY | Ce-2d | -93.9 | 8.49 | -100.5 | 8.43 | -100.0 | 8.30 | -100.0 | 8.13 |
| $^{D}$YDWPGDYCYLY | Ce-2d-Y3W | -83.8 | 6.06 | -93.3 | 5.92 | -92.9 | 5.88 | -95.0 | 5.85 |

**(B)**

| | | 2.5 min Incubation | | | | | |
|---|---|---|---|---|---|---|---|
| | | DAEMS Assay, Plate 1 | | DAEMS Assay, Plate 2 | | DAEMS Assay, Plate 3 | |
| Sequence | ID | Max I | pIC50 | Max I | pIC50 | Max I | pIC50 |
| $^{D}$YDYPGDYCYLYGTC-NHOH | Ce-1 NHOH | -99.2 | 9.94 | -101.1 | 9.93 | -101.2 | 9.96 |
| $^{D}$YDYPGDYCYLY | Ce-2d | -100.3 | 8.34 | -99.8 | 8.29 | -100.2 | 8.36 |
| $^{D}$YDWPGDYCYLY | Ce-2d-Y3W | -88.6 | 6.22 | -89.5 | 6.24 | -89.3 | 6.25 |

| | | 5 min Incubation | | | | | |
|---|---|---|---|---|---|---|---|
| | | DAEMS Assay, Plate 1 | | DAEMS Assay, Plate 2 | | DAEMS Assay, Plate 3 | |
| Sequence | ID | Max I | pIC50 | Max I | pIC50 | Max I | pIC50 |
| $^{D}$YDYPGDYCYLYGTC-NHOH | Ce-1 NHOH | -101.7 | 9.83 | -100.2 | 9.82 | -101.0 | 9.94 |
| $^{D}$YDYPGDYCYLY | Ce-2d | -99.3 | 8.04 | -99.6 | 8.09 | -99.5 | 8.14 |
| $^{D}$YDWPGDYCYLY | Ce-2d-Y3W | -82.2 | 6.17 | -88.9 | 6.12 | -87.2 | 6.23 |

| | | 10 min Incubation | | | | | |
|---|---|---|---|---|---|---|---|
| | | DAEMS Assay, Plate 1 | | DAEMS Assay, Plate 2 | | DAEMS Assay, Plate 3 | |
| Sequence | ID | Max I | pIC50 | Max I | pIC50 | Max I | pIC50 |
| $^{D}$YDYPGDYCYLYGTC-NHOH | Ce-1 NHOH | -100.0 | 9.64 | -100.0 | 9.62 | -101.0 | 9.54 |
| $^{D}$YDYPGDYCYLY | Ce-2d | -98.5 | 7.69 | -99.4 | 7.77 | -100.1 | 7.52 |
| $^{D}$YDWPGDYCYLY | Ce-2d-Y3W | -78.1 | 6.00 | -81.1 | 5.29 | -75.1 | 5.38 |

| | | 20 min Incubation | | | | | |
|---|---|---|---|---|---|---|---|
| | | DAEMS Assay, Plate 1 | | DAEMS Assay, Plate 2 | | DAEMS Assay, Plate 3 | |
| Sequence | ID | Max I | pIC50 | Max I | pIC50 | Max I | pIC50 |
| $^{D}$YDYPGDYCYLYGTC-NHOH | Ce-1 NHOH | -100.8 | 9.47 | -100.6 | 9.54 | -101.0 | 9.43 |
| $^{D}$YDYPGDYCYLY | Ce-2d | -99.9 | 7.37 | -99.9 | 7.52 | -100.3 | 7.39 |
| $^{D}$YDWPGDYCYLY | Ce-2d-Y3W | -64.8 | 4.60 | -65.6 | <5.00 | -67.9 | 4.97 |

**Supplementary Info 7** - Table of maximum inhibition and $pIC_{50}$ values calculated from the CRCs in Supplementary Info 7 for macrocyclic peptide inhibitors tested against 5 nM *C. elegans* iPGM enzyme across a 20 min time course for **(A)** the coupled-enzyme Kinase-Glo Plus (KGP) luminescence end point assay and **(B)** the DAEMS assay. Maximum inhibition values are the average of two duplicate wells treated with the highest concentration of compound per time point and $pIC_{50}$ values were calculated in GraphPad Prism with non-linear regression log((inhibitor) vs. response – variable slope (four parameters) with the constraint of absolute value Hill Slope less than 3, from the average of 2 replicate wells per concentration. Values were calculated for row-wise normalized data to no enzyme control as -100% inhibition per time point.

| Protocol Table . *C. elegans* co-factor independent phosphoglycerate mutase (iPGM) coupled-enzyme functional assay and DAEMS assay across 20 min time course. | | | |
|---|---|---|---|
| **Sequence** | **Parameter** | **Value** | **Description** |
| 1 | Peptides | 25 nL | Macrocyclic peptide inhbitors [stock solutions prepared at 5 mM based on mass]; 16-pt 1:3 titration series in duplicate across four replicate sets of columns for time course; or vehicle (DMSO); Compound transfer by Echo 555 acoustic dispense into 1536-well white/solid bottom, medium bind high base plate (Greiner) and Echo compatible cyclic olefin copolymer plate; plates froze at -80 $^{o}$C. |
| 2 | Reagent | 2 μL | **Enzyme solution:**<br>5 nM *Caenorhabditis elegans* (Ce) iPGM-HiBiT + Zn/Mn + Dialysis, final assay concentration, in standard 1X assay buffer<br><br>No enzyme control (column 1) and enzyme solution dispensed with BioRaptr 2 |
| 3 | Incubation | 20 min | Incubate enzyme + compounds at ambient temp for 20 min, protected from light |
| 4a | Reagent<br>(Coupled-Enzyme Assay) | 2 μL | Substrate solution; 0.4 mM final concentration 3-PG in coupled-enzyme substrate buffer; dispensed with BioRaptr 2 |
| 4b | Reagent<br>(DAEMS Assay) | 2 μL | Substrate solution; 0.4 mM final concentration 3-PG in 1X assay buffer; dispensed with BioRaptr 2 |
| 5 | Incubation | 2.5 – 20 min | Ambient temperature; dark for 2.5 – 20 min |
| 6 | Reagent | 1 μL | Ce-2d-K11 peptide quencher (added at 2.5, 5, 10 and 20 min post-substrate addition, 20 μM final concentration); dispensed with BioRaptr 2<br><br>DAEMS plate centrifuged at 1000 rpm for 2 min, sealed and frozen at -80 $^{o}$C until analysis |
| 7 | Reagent | 4 μL | Kinase-Glo Plus reagent (Promega) prepared according to manufacturer's protocol; dispensed with BioRaptr 2 to coupled-enzyme functional assay plate |
| 8 | Incubation | 10 min | Ambient temperature, protected from light |
| 9 | Detector | ViewLux | Read plate luminescence (Exposure = 1 sec; Gain = Medium; Speed = Slow; Binning = 2X) |
| **Step** | **Notes** | | |
| 2 | PGM enzyme buffer: 30mM Tris-HCl , pH 8.0, 5 mM $MgSO_4$, 20 mM KCl, 0.12% BSA + 7.5 nM enzyme.<br>• All solutions filtered through 0.22 μm syringe filter prior to dispense<br>• Ce_iPGM-C-HiBiT enzyme dialyzed with Zn/Mn (6.63 uM stock solution in 20% glycerol) | | |
| 4 | **Coupled-enzyme substrate solution**: 30mM Tris-HCl , pH 8.0, 5 mM $MgSO_4$, 20 mM KCl, 9 mM ADP, and 0.2 unit of enolase (Millipore Sigma Cat# E6126) and 0.3 unit pyruvate kinase (from rabbit muscle, Millipore Sigma Cat # P9136) + 800 μM 3PG (Millipore Sigma Cat #P8877).<br>**DAEMS substrate solution**: 30mM Tris-HCl , pH 8.0, 5 mM $MgSO_4$, 20 mM KCl + 800 μM 3-PG (Millipore Sigma Cat #P8877)<br>- Solutions filtered through 0.22 μm syringe filter prior to dispense | | |
| 6 | Ce-2d-K11 peptide quencher stock solution prepared at 10 mM in 1X PBS, pH 8.5; peptide diluted to 100 μM in diH20 for a final assay concentration of 20 μM in a salt concentration of 1.23 mM NaCl. | | |
| 7 | Kinase-Glo Plus reagent (Promega Cat # V3771) prepared according to manufacturer's recommendations and stored at -30 $^{o}$C until use.<br>- Reagent filtered through 0.22 μm syringe filter prior to dispense | | |

**Supplementary Info 8** – Assay protocol table for coupled-enzyme Kinase-Glo Plus (KGP) luminescence end point and DAEMS assays.

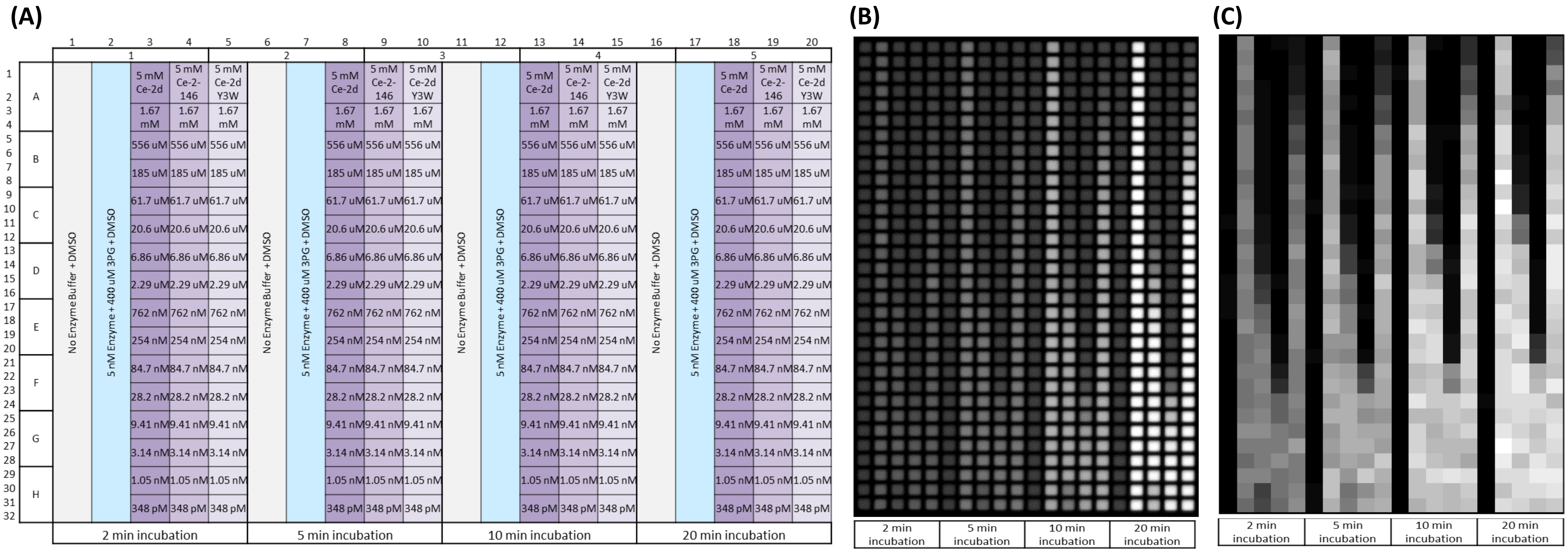


**Supplementary Info 9**- **(A)** Compound plate layout where peptides were titrated in a 16-pt, 1:3 dilution in duplicate wells down respective columns. Compounds were transferred to assay plates with an Echo 555 and diluted 160-fold in each assay. **(B)** Luminescence plate image from Perkin Elmer ViewLux CCD camera for the coupled-enzyme KGP luminescence functional assay imaged with the reader settings of 1 s exposure, medium gain, slow speed, 2x binning. Image is autocontrasted to highest and lowest luminescence signal where high signal is shown as lighter wells and low signal is shown as darker wells. **(C)** Simulated plate image using DAEMS data scaled to the lowest (black) and highest (white) signal intensities.

Isomers 2-phosphoglycerate (C3H7O7P · xLi+) and 3-phosphoglycerate ($C_3H_5Na_2O_7P$), DL-glyceraldehyde 3-phosphate ($C_3H_7O_6P$) and dihydroxyacetone phosphate ($C_3H_5O_6P \cdot 2Li$), chorismic acid ($C_{10}H_8O_6 \cdot Ba$) and prephenic acid ($C_{10}H_8O_6 \cdot Ba$), and citrate ($C_6H_5O_7Na_3.2H_2O$) were purchased from Sigma-Aldrich, and DL-threo-Isocitrate ($C_6H_5O_7Na_3.xH_2O$) was purchased from Cayman chemical. The macrocyclic peptide quencher Ce-2d K11 was purchased from GenScript and macrocyclic peptide inhibitors Ce-2d, Ce-1 NHOH and Ce-2d-Y3W were synthesized as previously described [1,2]. *C. elegans* iPGM was expressed and purified as described previously. [1]

All solvents used such as methanol, 2-propanol, ethyl acetate, and ammonium hydroxide were HPLC grade and purchased from Caledon laboratory chemicals.


1. Yu, H.; Dranchak, P.; Li, Z.; MacArthur, R.; Munson, M. S.; Mehzabeen, N.; Baird, N. J.; Battalie, K. P.; Ross, D.; Lovell, S., *Nature communications* **2017,** *8* (1), 14932.
2. 34. Wiedmann, M.; Dranchak, P. K.; Aitha, M.; Queme, B.; Collmus, C. D.; Kashipathy, M. M.; Kanter, L.; Lamy, L.; Rogers, J. M.; Tao, D., *Journal of Biological Chemistry* **2021,** *296*.


**Supplementary Info 10** – Reagents and Materials used.
